# Distinct antigenic properties of the SARS-CoV-2 Omicron lineages BA.4 and BA.5

**DOI:** 10.1101/2022.05.25.493397

**Authors:** Brian J. Willett, Ashwini Kurshan, Nazia Thakur, Joseph Newman, Maria Manali, Grace Tyson, Nicola Logan, Pablo R. Murcia, Luke B. Snell, Jonathan D. Edgeworth, Jie Zhou, Ksenia Sukhova, Gayatri Amirthalingam, Kevin Brown, Bryan Charleston, Michael H. Malim, Emma C. Thomson, Wendy S. Barclay, Dalan Bailey, Katie J. Doores, Thomas P. Peacock

## Abstract

Over the course of the pandemic variants have arisen at a steady rate. The most recent variants to emerge, BA.4 and BA.5, form part of the Omicron lineage and were first found in Southern Africa where they are driving the current wave of infection.

In this report, we perform an in-depth characterisation of the antigenicity of the BA.4/BA.5 Spike protein by comparing sera collected post-vaccination, post-BA.1 or BA.2 infection, or post breakthrough infection of vaccinated individuals with the Omicron variant. In addition, we assess sensitivity to neutralisation by commonly used therapeutic monoclonal antibodies.

We find sera collected post-vaccination have a similar ability to neutralise BA.1, BA.2 and BA.4/BA.5. In contrast, in the absence of vaccination, prior infection with BA.2 or, in particular, BA.1 results in an antibody response that neutralises BA.4/BA.5 poorly. Breakthrough infection with Omicron in vaccinees leads to a broad neutralising response against the new variants. The sensitivity of BA.4/BA.5 to neutralisation by therapeutic monoclonal antibodies was similar to that of BA.2.

These data suggest BA.4/BA.5 are antigenically distinct from BA.1 and, to a lesser extent, BA.2. The enhanced breadth of neutralisation observed following breakthrough infection with Omicron suggests that vaccination with heterologous or multivalent antigens may represent viable strategies for the development of cross-neutralising antibody responses.

## Main Text

In November 2021 a novel SARS-CoV-2 variant of concern, Omicron, was identified in Southern Africa^1^. The Omicron complex is now composed of five related lineages – BA.1 to BA.5, with BA.4 and BA.5 recently being described in Southern Africa^2^. The Omicron lineages BA.1 and BA.2 have been described as having a large antigenic distance from previous variants and current vaccine strains^3–5^, but a more modest antigenic distance between one another^6,7^.

As of May 2022, BA.4 and BA.5 have become the predominant variants in South Africa and are rising rapidly in several European countries. There are also clear signs BA.4 and BA.5 are driving a 5^th^ wave of cases in South Africa^2^. BA.4 and BA.5 encode identical Spike proteins and are most closely related to BA.2, containing several additional mutations –Δ69-70, L452R and F486V, but both are lacking the Q493R substitution relative to BA.2^2^. As L452R and F486V have been predicted to have an antigenic influence on Spike^8^, we examined the relative antigenic properties of the major Omicron complex family members, BA.1, BA.2, and BA.4 using human and animal vaccine sera, post-infection and vaccine-breakthrough sera, and therapeutic monoclonal antibodies (mAbs) in current use.

First we tested sera from triple-vaccinated individuals and found that these sera had a similar drop in neutralising titre for all Omicron lineages (6-15-fold), including an 8-to 10-fold drop against BA.4/BA.5 (Figure 1A). Using an older vaccinee cohort, we saw a similar pattern with similar drops to all Omicron lineages (Figure 1B,C). We used this same cohort to further investigate the effect of the 3^rd^ dose on BA.4/BA.5-specific titres. We found that for both three dose BNT162b2 and two dose ChAdOx1 + BNT162b2 boost vaccine regimes, the booster dose increased BA.4 neutralising titres by ≥10-fold, similar to that observed for BA.1 and BA.2.

**Figure 1.**
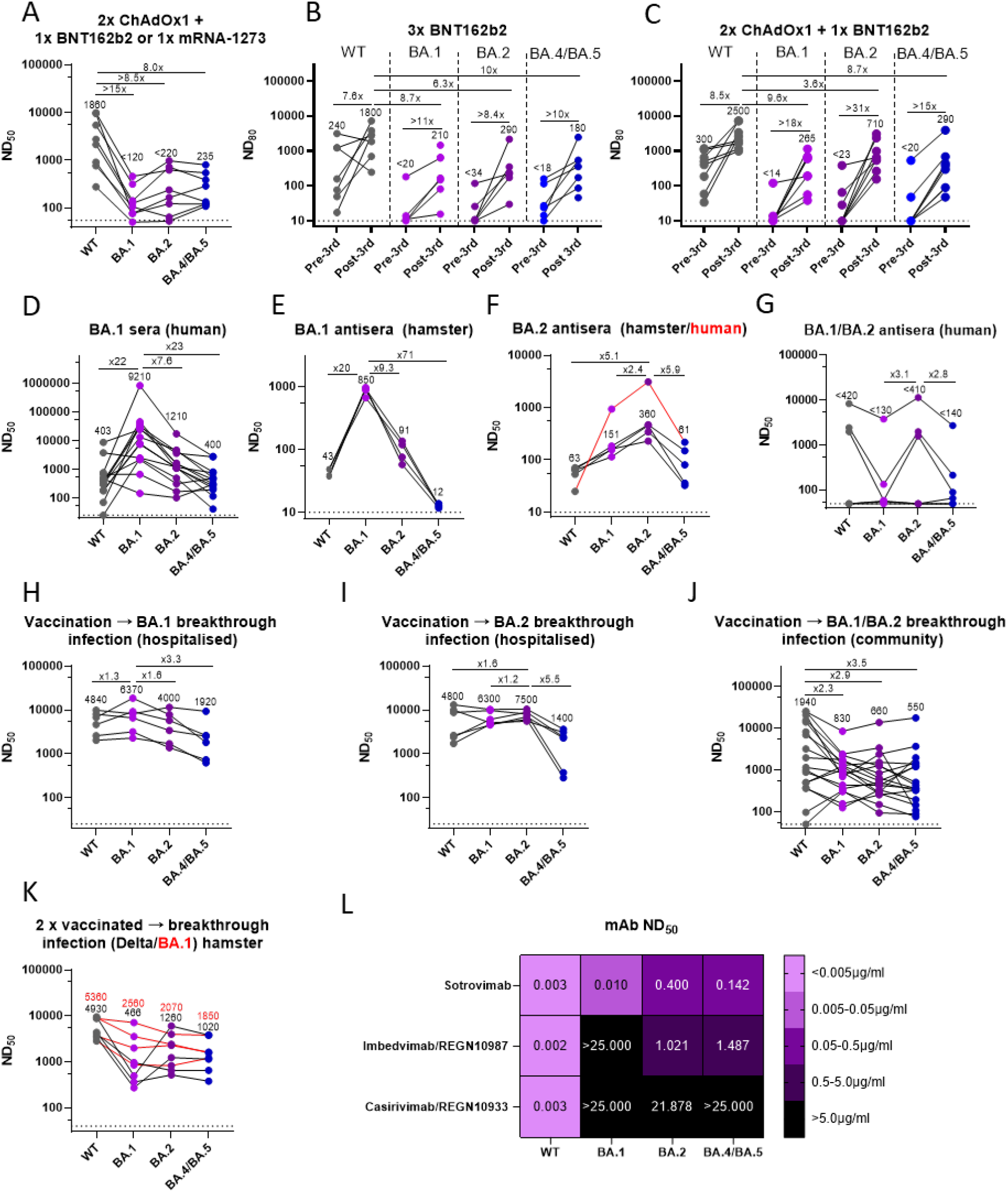
Antigenic properties of the BA.4/BA.5 Omicron Spike. A-C) Vaccine antisera from (A) the DOVE cohort vaccinated with two doses of ChAdOx1 (Oxford/AstraZeneca) with a booster dose of either BNT162b2 (Pfizer/BioNTech) or mRNA-1273 (Moderna) or (B,C) CONSENSUS geriatric cohort, vaccinated with two doses of ChAdOx1 or BNT162b2 and a booster dose of BNT162b2. Sera collected at the latest timepoint available prior to 3rd vaccine dose and at 4-6 weeks post 3rd dose. D-G) Post-infection sera from (D, F) hospitalised or (G) community infections. (E,F) Sera from previously infected hamsters. (H-J) Sera from Omicron-breakthrough infections of vaccinees in either (H,I) hospitalised or (J) the community. (K) Sera taken from breakthrough infections in hamsters vaccinated with two doses of an ancestral spike containing vaccine. (L) ND_50_ in μg/ml values for commonly used monoclonal antibodies against Omicron lineages. Neutralisation assays annotated with geometric mean titres and fold changes throughout.

We next investigated the cross-reactivity between different variants using human and hamster sera collected post-infection. Using sera from unvaccinated individuals with only a single known exposure to BA.1, we found a 23-fold drop in relative neutralising titres against BA.4/BA.5, and a more modest reduction against BA.2 (7.6-fold, Figure 1D). To validate the observations with human sera, which could potentially contain individuals who had asymptomatic exposure to pre-Omicron SARS-CoV-2 strains, we examined hamster sera collected post-infection (BA.1 or BA.2; Figure 1E,F). Consistent with the human sera, we found post-BA.1 hamster sera displayed a marked drop off in cross-neutralisation of BA.4/BA.5 (70-fold), while a more modest drop off was observed against BA.2 (9-fold). Hamster sera collected post-BA.2 infection displayed a less marked drop off in cross-neutralisation of BA.4/BA.5 (6-fold), while neutralisation of BA.1 was reduced by 2.3-fold. A similar pattern of cross-neutralisation was noted for a single human BA.2 infection-only sera (Figure 1F). We detected a similar pattern of cross -neutralisation with sera from unvaccinated people with unspecified Omicron infections. This cohort highlighted the extreme variability in immune responses to Omicron, with some individuals generating very poor post-infection antibody titres at the timepoints tested (Figure 1G). These data suggest that in naïve individuals, BA.4/BA.5 has a distinct antigenic profile, highly distinct to that of BA.1 and closer to that of BA.2.

We next tested sera from breakthrough infection, specifically from vaccinated individuals with known infections with BA.1 or BA.2, likely the most common exposure history in UK adults. Generally, Omicron breakthrough infections resulted in a robust pan-Omicron neutralising response. We observed reductions in neutralisation of 3.3-fold between BA.4/BA.5 and BA.1 for BA.1 breakthrough infection (Figure 1H) consistent with several other reports^9–12^, or 5.5-fold between BA.4/BA.5 and BA.2 for BA.2 breakthrough sera (Figure 1I). For community antisera from Omicron breakthrough infections where the lineage was unspecified, there were comparable drops in titre against all Omicron lineages (between 2.3 - 3.5-fold, Figure 1J). Further, we tested the responses of hamsters vaccinated twice with an ancestral vaccine strain and then infected with either Delta or BA.1. Surprisingly, we found Delta breakthrough after vaccination resulted in a cross-neutralising response to BA.2 and BA.4/BA.5 (< 2-fold drop), but not BA.1. BA.1 breakthrough in vaccinated hamsters also led to a moderately cross -neutralising response with BA.4/BA.5 being neutralised to a similar degree to BA.1 or BA.2 (<2-fold difference; Figure 1K). Overall, these data suggest that breakthrough infection with BA.1 or BA.2 post -vaccination leads to a cross-neutralising ‘pan-Omicron’ antibody response.

Finally, we tested the neutralisation activity of 3 commonly used mAbs – sotrovimab (a monotherapy developed by GSK), casirivimab/REGN10933 and imbedvimab/REGN10987 (which together make up the Regeneron cocktail, REGEN-COV/Ronapreve). We found that BA.4/BA.5 showed a broadly similar pattern of mAb sensitivity to that of BA.2 (Figure 1L), being recognised less well by sotrovimab than BA.1 or WT Spike, with marginally better recognition by imbedvimab than BA.1, as seen by others^10,12,13^.

To conclude, BA.4/BA.5 are antigenically distinct from previous Omicron variants, particularly BA.1, and are still at a substantial distance from the closely related BA.2. Sera elicited by breakthrough infections in vaccinated individuals with Omicron generally showed broad cross-neutralising activity, indicating that priming with an ancestral antigen and boosting with an Omicron-derived antigen (in this case BA.1 or BA.2 infection) elicits a pan-Omicron neutralising response. These data suggest that boost vaccination with Omicron-derived antigens could be an effective approach to inducing cross-protective immunity.

## Methods

### Neutralisation assays

Pseudovirus neutralisation assays were performed as described elsewhere^4,7,14–16^. Briefly, HEK 293T cells were transfected with lentiviral packaging plasmids and the named Spike construct - D614G(WT), BA.1, BA.2 or BA.4/5 to produce pseudovirus. Heat inactivated antisera was serially diluted and added to and mixed with a set volume of pseudovirus and incubated for 1 h at 37^o^C. Sera was then combined with target cells expressing human ACE2 (HEK293T or HeLa) and incubated for 48-72 hours. After this time cells were lysed and luciferase signals read by platereader. Antibody titre was then estimated by interpolating the point at which infectivity had been reduced to a set value of inhibition relative to the no serum control samples.

### Antisera and ethics

Hamster antisera was generated as we have described previously^7^, and was carried out under a United Kingdom Home Office License, P48DAD9B4.

For the first cohort of vaccine antisera, sera were collected from healthy volunteers participating in the COVID-19 **D**epl**o**yed **V**accin**e** Cohort Study (DOVE), or from the NHS Greater Glasgow and Clyde (NHSGGC) Biorepository. DOVE is a cross-sectional cohort study to determine the immunogenicity of deployed COVID-19 vaccines against evolving SARS-CoV-2 variants; a post-licensing cross-sectional cohort study of individuals vaccinated with deployed vaccines as part of the UK response to the COVID-19 pandemic. All DOVE participants gave informed consent to take part in the study, which was approved by the North-West Liverpool Central Research Ethics Committee (REC reference 21/NW/0073). Community sera were obtained from by NHSGGC Biorepository (ethical approval 550). Random residual biochemistry serum samples from primary (general practice) and secondary (hospital) healthcare settings were collected by the NHSGGC Biorepository from 2020 to 2022.

For the older adult vaccine cohort, healthy participants aged 50-90 years were recruited through North London primary care networks as part of the “COVID-19 vaccine responses after extended immunisation schedules” (CONSENSUS) study (UKHSA). Sera samples used in this study were taken from individuals vaccinated with 2 doses of Pfizer/BioNTech BNT162b2 (n=7) or 2 doses of Oxford/AstraZeneca ChAdOx1 nCoV-10/AZD1222 (n=8) 8-12 weeks apart, followed by a 3rd dose of Pfizer/BioNTech BNT162b2. Samples were analysed at the latest timepoint available prior to 3rd vaccine dose and at 4-6 weeks post 3rd dose. The protocol was approved by Public Health England Research Ethics Governance Group (reference NR0253; 27/01/21). Participants who were unable to provide informed written consent were excluded from the recruitment process.

For sequence confirmed BA.1 or BA.2 antisera surplus serum samples was approved by South Central – Hampshire B REC (20/SC/0310). SARS-CoV-2 cases were diagnosed by RT–PCR of respiratory samples at St Thomas’ Hospital, London. Vaccinated and BA.1 BTI – 5/6 had had 3 vaccine doses. Samples collected 18-27 days POS or post positive test. Vaccinated BA.2 BTI – 5/6 had had 3 vaccine doses. Samples collected 9-25 days POS or post positive test. SARS-CoV-2 naïve/BA.1 infected –18-27 days POS or post positive test. BA.1 or BA.2 variant infection were confirmed using whole genome sequencing as previously described ^14^ or using MT-PCR ^17^.

## Acknowledgements

We would like to acknowledge the UKHSA (formerly PHE) CONSENSUS team, as well as the vaccinated volunteers for their help and participation in supporting this study. This work was supported by the G2P-UK National Virology Consortium funded by the MRC (MR/W005611/1). Additional funding to D.B., N.T., and J.N. was by The Pirbright Institute’s BBSRC institute strategic program grant (BBS/E/I/00007038). The Medical Research Council (MRC) provided funding to E.T. for the COVID-19 DeplOyed VaccinE (DOVE) study (MCUU1201412) and to N.L., G.T., M.M. and B.W. (MC_UU_12014/1 & MR/W02067X/1). Health Data Research UK (HDR UK) funded E.T. and B.W. for the Evaluation of Variants Affecting Deployed COVID-19 Vaccine (EVADE) study (grant code: 2021.0155). K.D. and M.H.M. were supported by an award from the Huo Family Foundation.

## Author contributions

Data was collected by B.W. A.K., N.T., J.N., L.S., J.E., M.M., G.T., N.L., P.M., J.Z., K.S., G.A., K.B., and T.P. Data analysed by B.W., A.K., N.T., J.N., L.S., J.E., M.M., G.T., N.L., P.M., J.Z., K.S. and T.P. Data interpretation by B.W., D.B., K.D. and T.P. Writing and editing by B.W., D.B., and K.D. and T.P. funding acquisition by B.W. M.P., B.C., M.M., E.T., W.S.B, D.B., K.D. Supervision by B.W, W.B., D.B., and K.D. All authors were involved with editing and proofing prior to submission.

